# HIV-1 Mediated Cortical Actin Disruption Mirrors ARP2/3 Defects Found in Primary T Cell Immunodeficiencies

**DOI:** 10.1101/2023.07.27.550856

**Authors:** Jacqueline M. Crater, Daniel Dunn, Douglas F. Nixon, Robert L. Furler O’Brien

**Affiliations:** Department of Medicine, Division of Infectious Diseases, Weill Cornell Medicine, New York, NY, USA

**Author notes:** **Materials & Correspondence:** Correspondence and requests for materials should be addressed to Robert Furler O’Brien at.

**Keywords:** HIV-1, CD4^+^ T cell, Actin, Cytoskeleton, HIV-1 Nef protein, Cell morphology, Time-lapse imaging, HIV-1, ARP2/3, WAVE2, Wiskott-Aldrich syndrome, Primary Immunodeficiencies, Inborn Errors of Immunity, Actin, CD4^+^ T Cell Migration

## Abstract

During cell movement, cortical actin balances mechanical and osmotic forces to maintain cell function while providing the scaffold for cell shape. Migrating CD4^+^ T cells have a polarized structure with a leading edge containing dynamic branched and linear F-actin structures that bridge intracellular components to surface adhesion molecules. These actin structures are complemented with a microtubular network beaded with membrane bound organelles in the trailing uropod. Disruption of actin structures leads to dysregulated migration and changes in morphology of affected cells. In HIV-1 infection, CD4^+^ T cells have dysregulated movement. However, the precise mechanisms by which HIV-1 affects CD4^+^ T cell movement are unknown. Here, we show that HIV-1 infection of primary CD4^+^ T cells causes at least four progressive morphological differences as a result of virally induced cortical cytoskeleton disruption, shown by ultrastructural and time lapse imaging. Infection with a ΔNef virus partially abrogated the dysfunctional phenotype in infected cells and partially restored a wild-type shape. The pathological morphologies after HIV-1 infection phenocopy leukocytes which contain genetic determinants of specific T cell Inborn Errors of Immunity (IEI) or Primary Immunodeficiencies (PID) that affect the actin cytoskeleton. To identify potential actin regulatory pathways that may be linked to the morphological deformities, uninfected CD4^+^ T cell morphology was characterized following addition of small molecule chemical inhibitors. The ARP2/3 inhibitor CK-666 recapitulated three of the four abnormal morphologies we observed in HIV-1 infected cells. Restoring ARP2/3 function and cortical actin integrity in people living with HIV-1 infection is a new avenue of investigation to eradicate HIV-1 infected cells from the body.

---

Intracellular pathogens frequently manipulate the actin cytoskeleton at different stages in their replication cycles^1,2^. Cortical actin balances physical forces to provide an architectural scaffold that creates a barrier to the environment while dictating cell morphology, effector function, and migration. HIV-1, like other enveloped viruses, must surpass cortical actin barriers at both early entry and late budding stages^3^. Once infected, HIV-1 reduces the cellular levels of F-actin^4^ causing the cell to lose its polarization^5^. Furthermore, the infected cell has a reduced velocity^5,6^, aberrant directionality^6,7^, and a decreased capability of traversing endothelial barriers^5,6^ and migrating in dense complex environments^6^. Although HIV-1 induced morphological changes in primary and oncogenic CD4^+^ T cells on 2D surfaces was first reported to lose their polarization^7^, infected cells *in vivo* show prolonged cell polarization when migrating^7,8^. These discrepancies arise from cell type observed, environmental complexity of the system, and the time scale of observation.

Viral proteins including Nef and downstream effectors PAK2 and cofilin have been implicated in actin disruption with varied effects on leukocyte morphology and migration^9,10^. Although there is a general agreement that Nef reduces cell mobility, differential reports of Nef-induced changes in cell polarization and morphology exist between model systems. Although prolonged cell polarization is reportedly lost in two dimensional systems using T cell lines^7^, HIV-1 infected cells maintain polarization and have Env-mediated elongated uropods in animal models and three dimensional culture systems^7,8^. The use of primary versus oncogenic cell lines, the environment complexity, and the time scale of observation may all be variables that lead to these discrepancies. To address these differences of HIV-1 and Nef-mediated alterations on primary T cell morphology, we combined ultrastructural microscopy with time-lapse imaging of HIV-1 infected cells. We observed the previously reported morphological changes that occur following HIV-1 infection and report additional phenotypes that provides insight into the molecular underpinnings of the general immunodeficiency that arises during progressive Acquired Immunodeficiency Syndrome (AIDS).

Here, we unify these disparate findings with time-lapse and ultrastructural microscopy showing that HIV-1 infection of primary CD4^+^ T cells causes at least five progressive morphological differences as a result of virally induced cortical cytoskeleton disruption. Infection with a Nef-deleted (ΔNef) virus partially abrogated the dysfunctional phenotype in infected cells. Along with a recently reported lamellipodial blebbing phenotype^7^, we observe a fifth unreported pathological morphology distinct to HIV-1 infected CD4^+^ T cells. These two morphologies phenocopy migrating cells which contain genetic determinants of specific T cell primary immunodeficiencies (PIDs) also known as Inborn Errors of Immunity (IEI) that affect the actin cytoskeleton, particularly diseases affecting the ARP2/3 complex. The branched actin nucleator ARP2/3 is an important component of the lamellipodia during migration as well as the T cell immune synapse during activation and is frequently targeted by intracellular bacteria and enveloped viruses^1,2^.

Our study qualitatively characterizes cortical actin changes in migrating CD4^+^ T cells that give rise to unreported morphologies in HIV-1 infected cells. Furthermore, we show that the actin disruption during HIV-1 replication is reminiscent of actinopathies that give rise to several PIDs/IEI. These aberrant morphologies phenocopy leukocytes with defective ARP2/3 branching found in certain PIDs/IEI. In these genetic disorders, ARP2/3 dysfunction can directly lead to T cell depletion, dysfunction, or systemic localization because of ARP2/3’s role in chemotaxis ^11-13^ and immune synapse formation^14-22^. Furthermore, the ARP2/3 complex is manipulated by several intracellular pathogens including bacteria like *Rickettsia, Listeria, and Shigella*^2^ and viruses like *Vaccinia* and retroviruses^1^. The ARP2/3 complex’s pivotal role in pathogenic infection and systemic immunodeficiencies makes it a lucrative target for HIV-1 proteins that disrupt the actin cytoskeleton. The mechanically destabilized cellular cortex found in leukocytes from PIDs/IEI and following HIV-1 infection may provide context into cellular and systemic pathologies that drive similar clinical manifestations observed in primary and acquired immunodeficiencies.

## Results

### HIV-1 Infected CD4^+^ T Cells Exhibit Cortical Actin Disruption that Lead to Aberrant Morphologies

HIV-1 infection is known to cause actin dysregulation and alter chemotaxis in CD4^+^ T cells, primarily due to the effects of the Nef protein^23-25^. Infected cells have significantly less F-actin^4^, lose their polarization^5^, have reduced velocity^5,6^ and directionality^6,7^, and lose their ability to traverse endothelial barriers^5,6^ and migrate in more complex dense environments^6^. Although HIV-1 induced morphological changes in primary and oncogenic CD4^+^ T cells on 2D surfaces are reported to lose their polarization^7^, infected cell migration *in vivo* show prolonged polarization with Env-mediated elongated uropods and occasional polarized lamellipodial blebbing in 3D collagen matrix models^7,8^. These disparate results are likely due to the environment complexity, image resolution, and time scale in which these experiments were observed. To further characterize morphological changes within HIV-1 infected primary human CD4^+^ T cells at higher resolution, ultrastructural microscopy was combined with 2D time-lapse imaging at shorter time scales.

Negatively selected CD4^+^ T cells from 11 normal progressors and 15 healthy donors were activated and expanded prior to infection. Although cells were infected with either patient-derived or laboratory-adapted HIV-1 isolates, similar morphological abnormalities were present during time-lapse imaging regardless of viral isolate used. Transmission electron microscopy (TEM) was done on these samples and five distinct structural abnormalities were characterized.

Uninfected migratory CD4^+^ T cells have a polarized morphology with a leading edge or lamellipodium containing dynamic branched and linear F-actin structures that bridge intracellular components to surface adhesion molecules. These actin structures are tethered to the nucleus, which is anterior to and connected to microtubule and intermediated filament networks beaded with membrane-bound organelles in the trailing uropod (**Figure 1 and Supplementary Video 1)**. Along with migratory cells, there are typically abundant rounded non-migratory cells in the absence of chemokine addition. These non-polarized cells can easily be seen in both infected and mock conditions (data not shown) and is indicative of the non-polarized infected cells previously reported^5^. Two additional structural abnormalities we observed in HIV-1 infected cells, rounded cell blebbing (**Figure 2A and Supplementary Video 2**) and multinuclear syncytia (**Supplementary Video 3**), have previously been reported by multiple groups.

**Figure 1:**
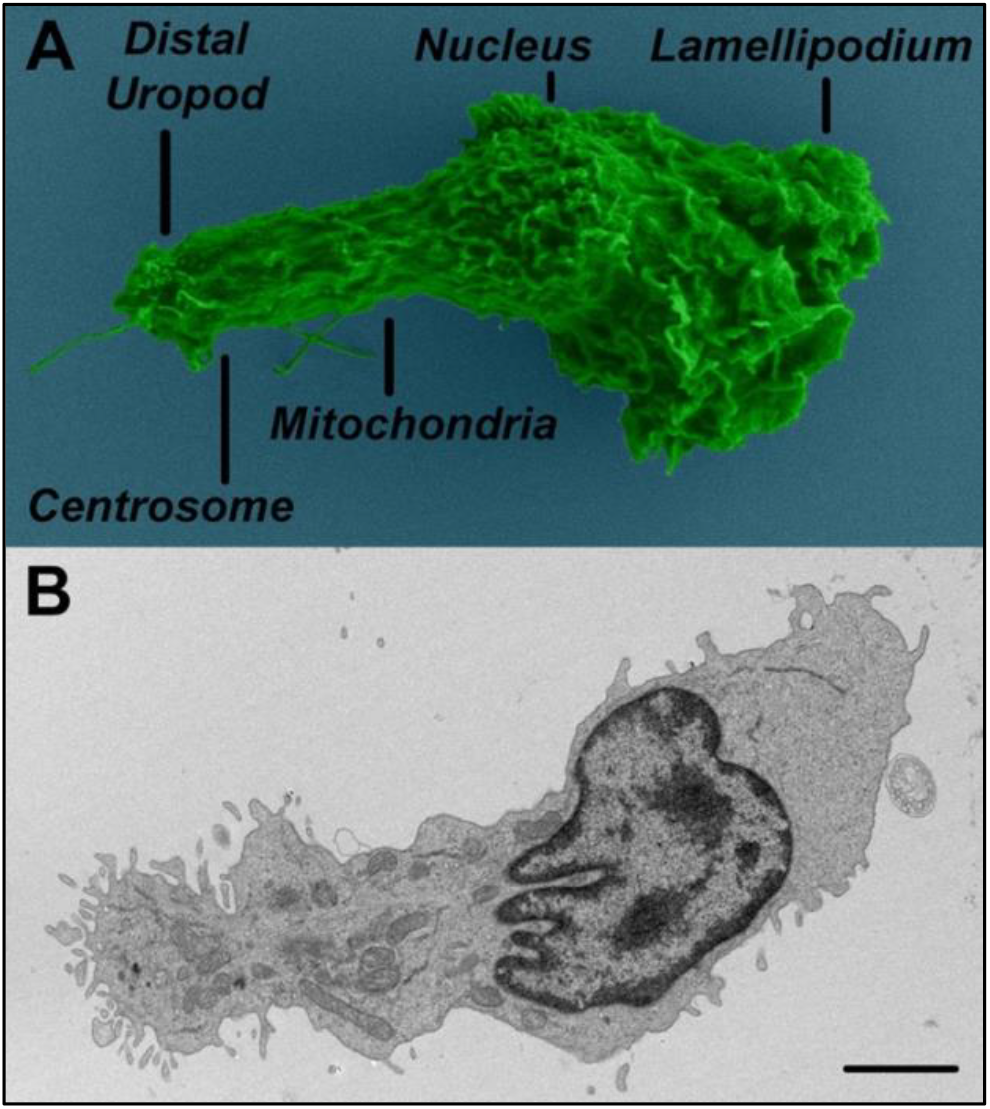
Ultrastructure of uninfected primary human CD4^+^ T cells migrating on fibronectin. As seen in the scanning electron micrograph (A), the lamellipodium is at the leading edge of the migrating cell while the distal uropod trails at the rear. (B) An anatomical view of organelles and cytoskeleton is seen in the transmission electron micrograph. Scale = 2μm

**Figure 2:**
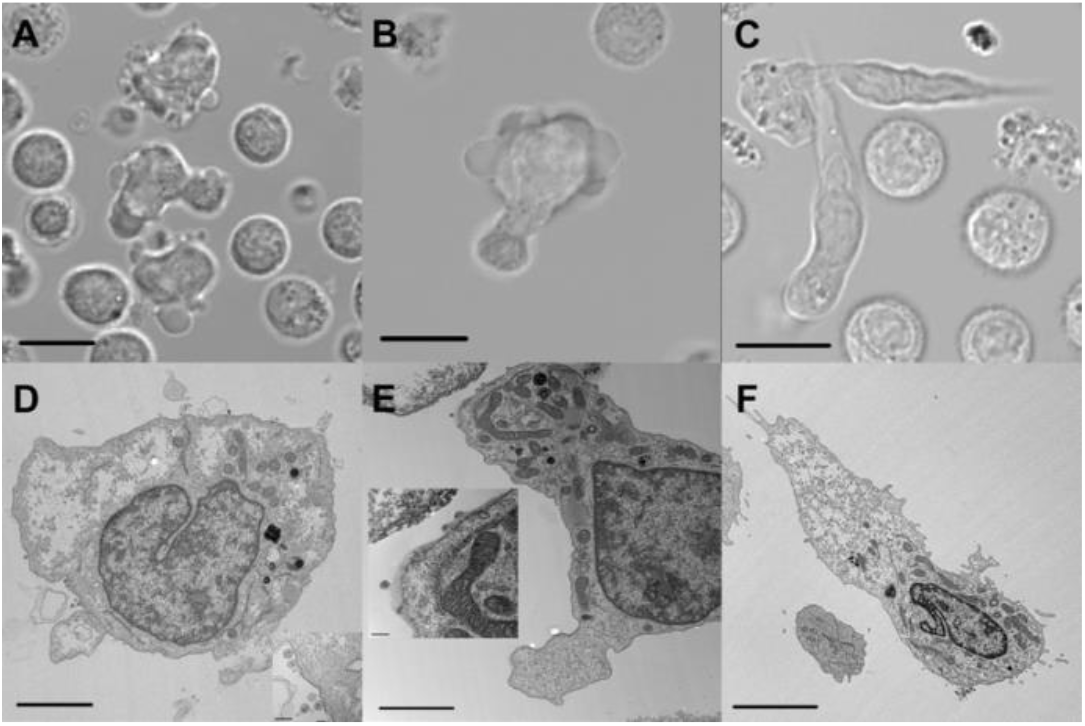
In addition to syncytia, HIV-1 infection leads to at least four distinct morphological differences in primary human CD4+ T cells. Apolarized rounded cells previously reported can be found with cells that have (A) rounded blebbing (B) polarized lamellipodial blebbing (C) or a needle-like or “Rhino” phenotype. Scale = 10μm. Organelle locations, actin disruption, and virion production for these morphologies can be seen in TEM micrographs respectively

The fourth abnormal phenotype of HIV-1 infected CD4^+^ T cells we observed was non-apoptotic blebbing polarized at the lamellipodia. These cells have typically extended uropods and clearly produce virions at the distal uropod by TEM (**Figure 2B, 2E, and Supplementary Video 4**). Although this phenotype was observed in 2-dimensional cultures during this study, similar cortical actin disruption was recently reported in HIV-1 infected T cells migrating in 3-dimensional collagen matrices^7^.

In addition to the previously reported morphologies, we observed a new abnormality that provides insight into the underlying cellular pathology induced by HIV-1. This final morphological abnormality is an elongated and pointed lamellum/lamellipodium sometimes but not exclusively with a retracted uropod (**Figure 2C, 2F, and Supplementary Video 5**). To distinguish the lamellipodial structure from the uropod, vital dyes recognizing the nucleus and mitochondria were added. The elongated structure was void of membrane-bound organelles that localize to the uropod, indicating the abnormal extension is indeed the leading edge of the cell. The nucleus and granular organelle-rich uropod can be visibly distinguished from this structure even under brightfield conditions. This phenotype is an abnormal pointed elongation of the lamellum/lamellipodium and should not be confused with uropod retraction defects previously shown to be HIV-1 Env-dependent in lymphoid tissues^8^.

These five abnormal phenotypes were consistently observed with both R5 and R5/X4 tropic viruses ADA-M, HIV-1 92/BR/014, and HIV-1 92/BR/004. To discern if these abnormal morphologies were unique to HIV-1 infected cells or present in uninfected bystander cells, a fully intact R5-tropic virus containing both the *nef* gene and an IRES-EGFP reporter^7^ was used to infect cells prior to time-lapse imaging. As shown in **Figure 3 and Supplementary Video 6**, these cortical actin abnormalities were unique to the infected (EGFP^+^) cells. Neighboring uninfected cells migrate normally with unaltered morphologies, suggesting an intraceullular viral protein is key to the observed cortical actin disruptions.

**Figure 3:**
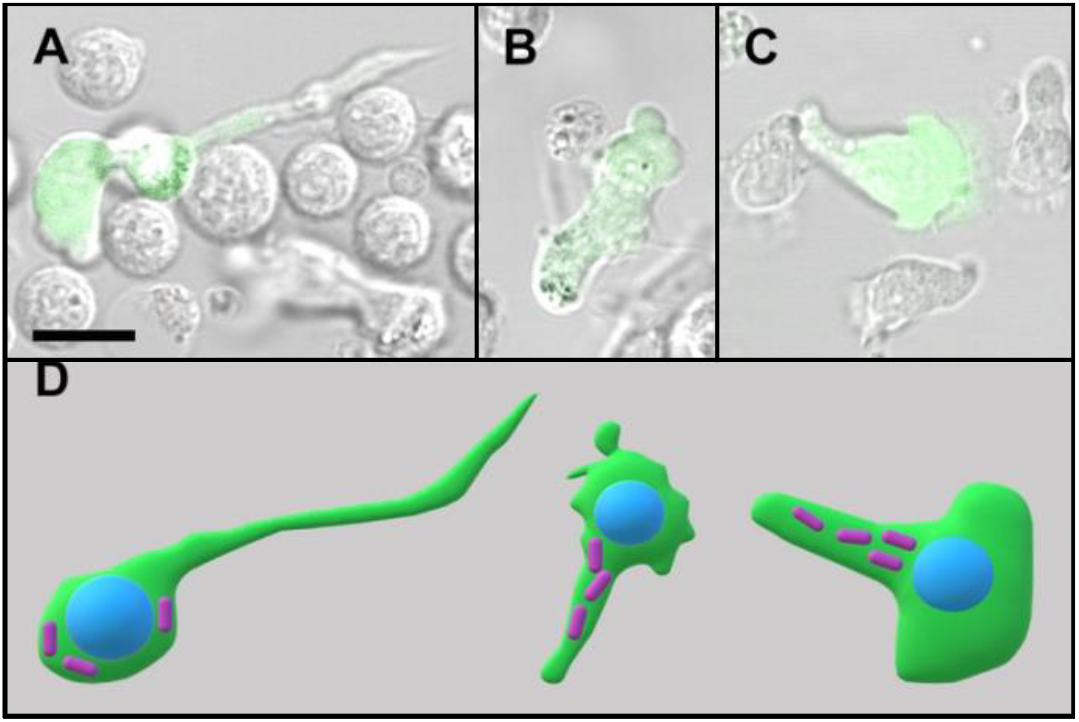
Morphological changes occur in HIV-1 infected cells and not bystanders. (A) The Rhino and (B) polarized lamellipodial blebbing only occur in cells infected with a fully intact HIV-1 EGFP reporter virus, not in bystander cells. (C) However, the absence of Nef results in a partially restored morphology with typical broad lamellipodia and uropod extension; Scale = 10μm. (D) Cartoons of morphologies with respective location of nuclei (blue) and mitochondria (fuchsia rods) shown for anatomical orientation.

### HIV-1 Nef is Not the Sole Viral Factor that Destabilizes Cortical Actin

The influential role of lentiviral Nef proteins in disease progression began to be uncovered in the early 1990s following several clinical reports and *in vivo* animal studies that interrogated Nef function^26-29^. Although multiple HIV-1 proteins are linked to actin dysregulation^3^, membrane localized Nef has repeatedly been shown to alter cortical actin *in vitro* and *in vivo*^4-8,23-25^. Several of these studies indicate that the actin disruption and subsequent polarization and migration defects were caused by the Nef associations with PAK2 and the Nef-associated kinase complex^5,7^.

To confirm the role of Nef in the aberrant morphologies we observed in our study, we characterized infected cell morphology following infection with wild-type (WT) or *nef*-deleted (ΔNef) IRES-EGFP reporter viruses. Although infection with ΔNef almost completely restored wild-type polarization and lamellipodial structure, non-apoptotic blebbing still occurred, albeit at a greatly decreased frequency (**Figure 3C and Supplementary Video 7)**.

This suggests that although HIV-1 Nef is a major disruptor of cortical actin, there may be additional viral mechanisms outside of Nef that induce actin instability. This is consistent with previous findings that depending on the donor, Nef deletion partially or completely restores T cell polarization and additional viral factors may influence T cell morphology and migration^5^. Although donor genetics may influence Nef’s ability to disrupt cortical actin, other groups have looked at the role of additional viral proteins. HIV-1 Vpu was a likely candidate since it has several overlapping roles with Nef; however, others have shown that Vpu does not impact cell morphology or migration speed in absence of Nef^7^. Other groups have recently linked HIV-1 Gag to actin disruption^30,31^, however, the use of cell lines and non-T cells in these studies makes it challenging to interpret. Gag is an interesting candidate since both HIV-1 Nef and Gag are myristoylated and closely interact with actin at the plasma membrane^3^. Although intrinsic viral redundancy in actin disruption may exist, Nef is likely the dominant effector and its absence is enough to delay or inhibit disease progression^26,32,33^.

### Actin Disruption in HIV-1 Infected Cells Mirrors ARP2/3 Inhibition

Aside from the apolarized, rounded blebbing, and syncytial phenotypes, the two newly reported morphologies may give insight into HIV-1 induced cellular pathology and subsequent disease progression. These unique morphological changes mimic migrating leukocytes from PIDs/IEI caused by mutations in actin regulatory genes. Elongated phenotypes are found in migrating leukocytes following depletion of DOCK8, CDC42, and PAK1/2^34^; however, these leukocytes have drastically elongated nuclei which we did not observe in HIV-1 infected CD4^+^ T cells. Inhibiting many of these actin regulatory proteins, including CDC42, Rac1/1b/2/3, and PAK2, induced varied phenotypes that do not resemble HIV-1 infected cells (**Supplemental Video 8)**.

The first aberrant morphology we observed in our 2D migration studies was polarized lamellipodial blebbing, similar to a phenotype recently reported by another group using 3D collagen matrices^7^. Polarized blebbing migration has been reported in several cell types outside of HIV-1 infection (**Table 1**), with a unifying mechanism surrounding ARP2/3 complex inhibition^35-42^. ARP2/3 is important in nucleating cortical actin branching at the lamellipodia of migrating T cells as well as organizing the immune synapse during T cell activation.

**Table 1.**
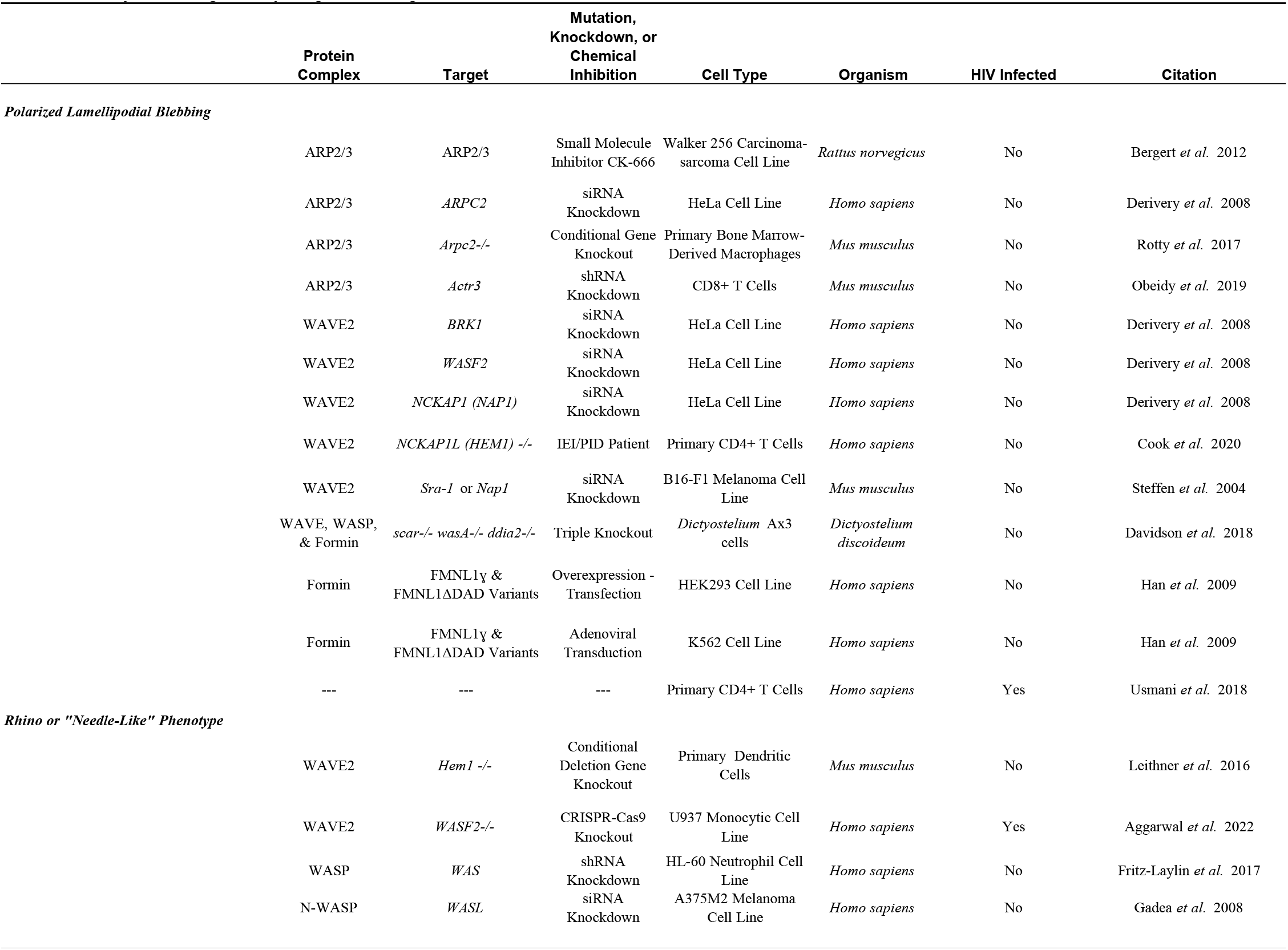
Summary of Pathological Morphologies Following ARP2/3 Inhibition

The previously unreported fifth phenotype we observed – a pointed elongated lamellipodia - provides further evidence that HIV-1 inhibits the ARP2/3 complex. This elongated phenotype was first described in leukocytes that were deficient in the WAS protein, which the authors coined “Rhino” shaped^43-45^. In addition to WAS knockdowns, inhibiting the related WAVE2 complex also induces this fifth phenotype. Our imaging of HIV-1 infected T cells is strikingly similar to the WAVE2-deficient leukocytes reported by the Sixt group^45^. Their time-lapse and TIRF imaging of the WAVE2-deficient cells is complemented with SEM and electron tomography which unearths the branched actin filament disruption observed in these “needle-like” cells. A summary of reports describing this morphology in the literature is also found in **Table 1**.

Previous reports indicate that membrane blebbing in other cell types can be induced by disrupting the branched cortical actin structures in human cells through chemical inhibition of the ARP2/3 complex with the small molecule inhibitor CK-666^35,46,47^. Since the phenotypes of ARP2/3-inhibited cells from other studies strongly mirror the aberrant morphologies we observed in HIV-1 infected cells, we characterized the morphology of primary human CD4^+^ T cells following chemical inhibition of the ARP2/3 complex with CK-666. Following addition of the small molecule inhibitor of ARP2/3, CK-666, both “Rhino” and polarized lamellipodial blebbing phenotypes were observed in uninfected CD4^+^ T cells (**Figure 4 and Supplementary Video 9)** whereas the non-active small molecule analog CK-689 had no observable effect. The presence of two aberrant morphologies we see in primary CD4^+^ T cells following ARP2/3 inhibition is consistent with the Sixt group’s findings of two TLR-driven differentiation-dependent morphologies they observed in dendritic cells following knocking out WAVE2^45^. This suggests that the lamellipodial blebbing and the needle-like phenotype we see in HIV-1 infection may be dependent on the maturation status of the CD4^+^ T cell. Our initial findings indicate that IL-18 differentiates cells to the needle-like or “Rhino” phenotype (data not shown), suggesting a maturational link between these two populations.

**Figure 4:**
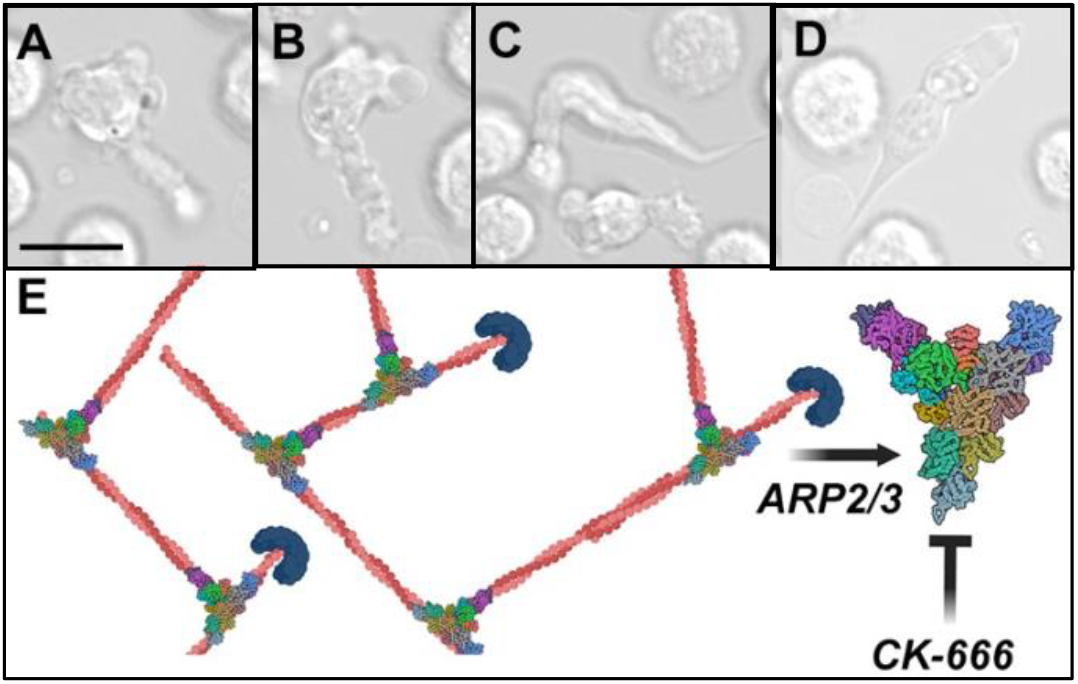
Chemical inhibition of ARP2/3 in uninfected CD4^+^ T cells recapitulates morphological differences seen in HIV-1 infected cells. Polarized lamellipodial blebbing (A-B) and Rhino morphologies (C-D) occur following addition ARP2/3 small molecule inhibitor CK-666, Scale = 10μm. (E) ARP2/3 is a multi-protein complex that leads to dendritic branching of the actin cytoskeleton. PDB ID: 7AQK

In order to measure differential changes in ARP2/3 components or its regulators following HIV-1 infection, we used mass spectrometry to quantify label-free proteins that were differentially expressed in sorted HIV-1 WT, HIV-1 ΔNef, and mock infected cells. A total of 5559 proteins were identified, including six HIV-1 proteins and EGFP in ten biological replicates **(Extended Data Table 1)**. Although housekeeping proteins like glycolytic enzymes (GAPDH) and cytoskeleton proteins (actin) are commonly used as proteomic normalization controls, these proteins are differentially expressed in T cells depending on activation status. To compare the 3 different populations of cells (HIV-1 WT, HIV-1 ΔNef, and HIV-1 negative), live singlet EGFP^+^ cells were sorted and isolated protein reads were normalized to the centrosomal-localized protein γ-tubulin. We did not detect any significant alteration in ARP2/3 complex proteins or known regulatory proteins within infected cells. This suggests that if ARP2/3 inhibition is occurring, it may not be due to degradation of ARP2/3 complex members. Numerous post-translational modifications and inhibitory proteins of these complexes have been reported to affect their activity or sequestration within different intracellular locations^48-50^. The role of cofilin, PAK2 or other Nef-associated enzymes in possible post-translational modifications of ARP2/3 or its regulators needs to be further examined.

## Discussion

A key finding in this study is that polarized lamellipodial blebbing and “Rhino” or “needle-like” morphological phenotypes are specific to productively infected CD4^+^ T cells from normal progressors and all healthy controls. Elongated pointed lamellipodia and non-apoptotic polarized blebbing, similar to the phenotypes of HIV-1 infected cells we observed in this study, occur following mutations in actin regulatory proteins^34^. These abnormalities are observed regardless of infection with various HIV-1 reporter and primary viral isolates and are predominantly but not exclusively due to Nef. HIV-1 infected cells strikingly mirror cells with defective ARP2/3-mediated actin branching. Cells with inhibited ARP2/3 activity, whether induce by small molecule inhibitors, gene knock-downs, or presence of negative protein regulators, exhibit both needle-like and polarized lamellipodia blebbing phenotypes^43-45^. Although direct evidence of HIV-1 induced ARP2/3 inhibition is not shown in this study, there are multiple lines of evidence that warrant further investigation into the role ARP2/3 dysfunction in the cellular pathology induced by HIV-1.

Potential mechanisms of HIV-1 induced ARP2/3 are still under investigation. Although ARP2/3 functionality was previously shown to be required for the early steps of HIV-1 infection^1,51^, others have indicated that ARP2/3 functionality may be altered at the budding stages later in the replication cycle^7,52,53^. Both Nef and Gag are membrane-localized N-myristoylated proteins and may work synergistically to inhibit ARP2/3 in infected cells. HIV-1 Nef has been linked to N-Wasp dysfunction in cell lines^31^ while HIV-1 Gag has been linked to ARP2/3 activity by others^30,31^. Activation of the WAVE2-IRSp53(BAIAP2)-ARP2/3 cascade was previously implicated in viral budding in the late stages of viral replication^30^, however these studies were done in Jurkat T cell lines and we did not detect the IRSp53(BAIAP2) protein in our proteomics studies of primary human CD4^+^ T cells. Furthermore, this same group has recently submitted a preprint suggesting that HIV-1 Gag inhibits rather than activates ARP2/3 through an interaction with the ARP2/3 inhibitor Arpin, again in cell lines. Although the Arpin protein was not present in our proteomics screen of primary CD4^+^ T cells, the related AP3S2-Arpin fusion protein was present in low levels. The known interactions between HIV-1 Gag and the AP3 complex makes this a compelling hypothesis; however, additional analysis will be needed to confirm the role of a Gag-Arpin mediated inhibition of ARP2/3 in primary CD4^+^ T cells. Although the role of IRSp53^4,30^, Arpin, and N-Wasp^31^ have been implicated in HIV-1 Gag and Nef mediated actin disruption, none of these proteins were present in our proteomic screen of primary human CD4^+^ T cells and direct evaluation of HIV-1 mediated ARP2/3 inhibition in primary CD4^+^ T cells still needs confirmation. The roles of cofilin, PAK2 or other Nef-associated enzymes in possible post-translational modifications of ARP2/3 or its regulators also needs to be confirmed.

Patients with primary immunodeficiencies that alter ARP2/3 activity also have decreased T cell absolute numbers and function and are more susceptible to opportunistic infections, similar to the clinical characteristics of AIDS. In 1937, Wiskott reported on an inherited immune disorder affecting males in a family^54^ and later in 1954, Aldrich *et al*. reported on a family exhibiting a similar sex-linked recessive disease affecting males in a pedigree who were anemic, had bloody diarrhea, and had repeated opportunistic infections including oral thrush^55^. Over the next two decades, additional cases were reported in the literature showing similar immunologic impairments primarily in the thymus-dependent compartment of cellular immunity. This grouping of cases was diagnosed as a new disease named after the two physicians who contributed to the initial reports – Wiskott-Aldrich Syndrome (WAS)^56^. Following WAS, several additional actin-regulatory genes that affect the ARP2/3 complex

(**Figure 5**) have been implicated in primary immunodeficiencies that affect T cell function including *ARPC1B*^57,58^, *ARPC5*^*59*^, *WIPF1, CORO1A, DOCK2, DOCK8, DOCK11, CDC42, RAC1*, and *RAC2*^*60*^ among others^61^. Genetic defects in the *NCKAP1L (hem1)* gene, which the Sixt group deleted to make WAVE-deficient leukocytes^45^, was also recently identified in human patients that exhibit a novel immunodeficiency^62,63^ with recurrent infections and increased autoimmunity. Several murine models with conditional knockouts of the WAVE2 complex show immunological characteristics similar to AIDS including inverted CD4/CD8 ratios, skewing of T cells toward exhausted memory phenotypes, and enhanced autoimmunities^45,62,64^.

**Figure 5:**
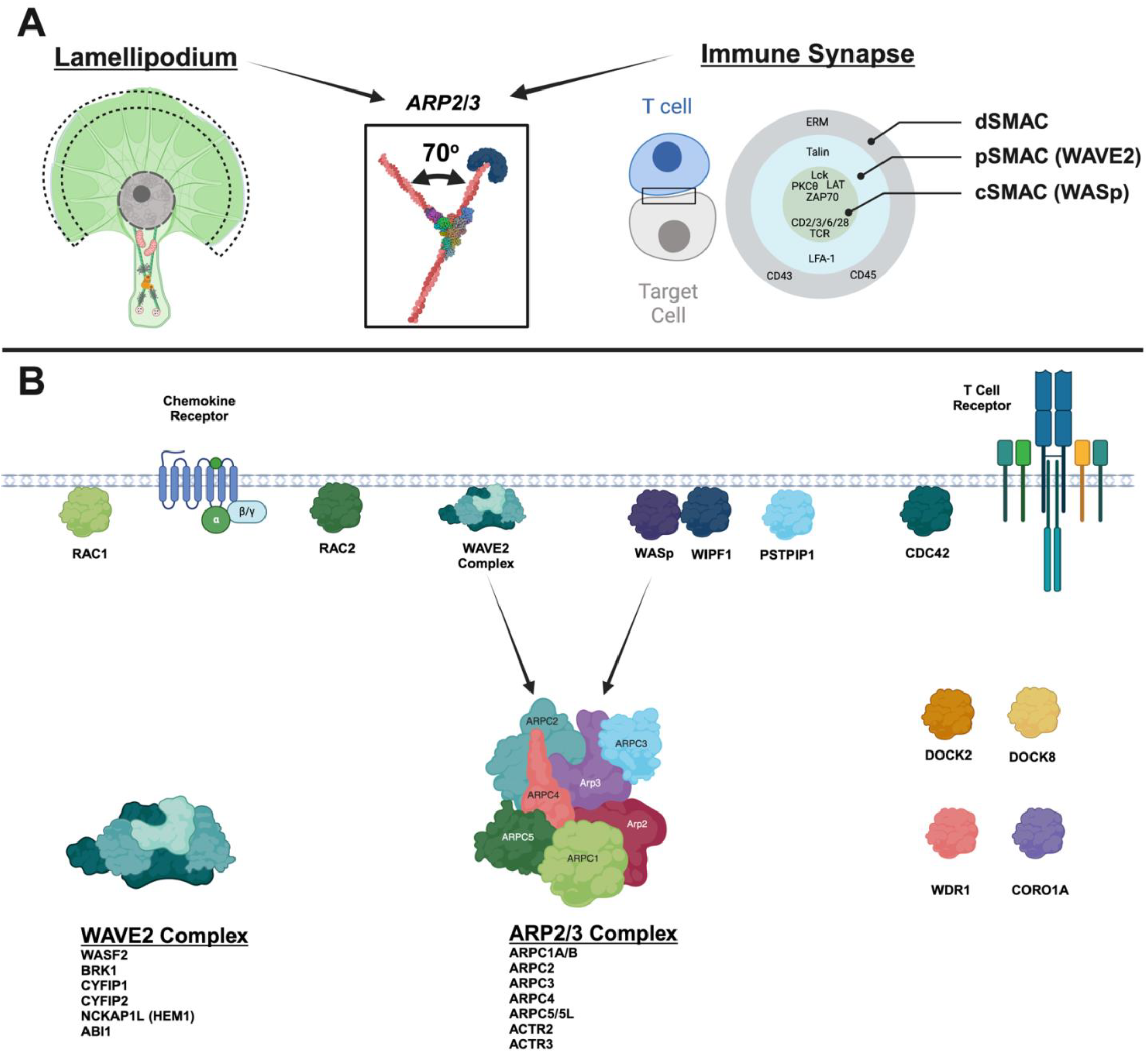
The branched actin nucleator ARP2/3 is central to leukocyte function and when inhibited causes Primary Immunodeficiencies (PIDs) or Inborn Errors of Immunity (IEI). (A) Among other intracellular locations, ARP2/3 functions at the lamellipodium of migrating leukocytes as well as the immune synapse during cell activation. (B) Mutations within ARP2/3 complex components or upstream regulators of ARP2/3 are known determinants of several congenital diseases that lead to immunodeficiencies.

The loss of ARP2/3 functionality in PIDs/IEI along with the known hijacking of the ARP2/3 complex by intracellular pathogens emphasizes the pivotal role of ARP2/3 in immunity. The growing body of evidence that viral proteins including HIV-1 Nef and potentially Gag inhibit ARP2/3 activity may provide additional clues to how HIV-1 mediated actin disruption causes cellular pathology and systemic immunodeficiency.

## Online Methods

### CD4^+^ T cell isolation and activation

Peripheral blood samples from deidentified participants were obtained from the Gulf Coast Blood Center (Houston, Texas, USA). All participants signed informed consent forms approved by their respective Investigational Review Boards (IRB). Freshly acquired blood was processed by Ficoll-Paque separation to isolate peripheral blood mononuclear cells (PBMC) from healthy donors and normal HIV-1 progressors. Negative selection of total CD4^+^ T cells was done using StemCell EasySep Human CD4^+^ T Cell Enrichment Kit (Catalog #19052). Cells were than cultured in RPMI, 10% fetal bovine serum (FBS), penicillin, streptomycin, and glutamine at one to two million cells/mL. Infections were done on negatively selected total human CD4^+^ T cells that were previously activated with anti-CD3 (Ultra-LEAF™ Purified anti-human CD3 Antibody, clone OKT3, Biolegend, Catalog #317325) and anti-CD28 (Ultra-LEAF™ Purified anti-human CD28 Antibody, Clone CD28.2, Biolegend, Catalog #302933) antibodies in the presence of rhIL-2 and IL-15. The media was changed every three to four days for two weeks prior to infection. Post-activation and prior to infection, a second round of negative selection was done to achieve 97-99% CD4^+^ T cell purity as measured by flow cytometry. The cells were then resuspended at a concentration of 2*10^7^ cells/mL in RPMI supplemented with 10% FBS, penicillin, streptomycin, glutamine, IL-2 and IL-15 prior to infection.

### Infection

For each of the infection and mock conditions, 500uL of cells were added to a well in a 6-well plate before adding 500uL of virus or mock supernatant. The cell/virus mixture was spinoculated at 2900 rpm (1965g) for 2 hours at 37°C. These cells were added to an equal number of non-spinoculated cells and cultured at 2*10^6^ cells/mL at 37°C for up to 3 weeks, with microscopic analysis and media changes occurring every 2-3 days. To optionally increase the population of infected cells, cells were crowded at Day 3 post-infection by culturing in 200uL at 2*10^6^ cells/mL in 96-well ‘U’-bottom plates for an additional 3 to 4 days. Monitoring of infected cells populations was done by intracellular HIV-1 Gag p24 measurements using flow cytometry and measuring EGFP fluorescence by microscopy when using reporter viruses.

Cells were infected with primary, laboratory, and reporter isolates of CCR5-tropic or CCR5/CXCR4 dual-tropic Human Immunodeficiency Virus-1. The following three viruses were obtained from the NIH AIDS Reagents program: HIV-1 ADA (ARP-416, CCR5-tropic), HIV-1 92/BR/014 (ARP-1753, Dual-tropic, syncytium inducing), HIV-1 92/BR/004, ARP-1752 (CCR5-tropic, non-syncytium inducing). Reporter viruses were generated from the following plasmids that were generously provided by Dr. Thorsten Mempel and Dr. Shariq Usmani: HIV-1 (WT) IRES-EGFP (pBR43IeG nef+ R-BaL env) and HIV-1 (ΔNef) IRES-EGFP (pBR43IeG nef+ R-BaL env, Nef deleted)^7^. Plasmids were transformed into Stbl2 competent *E. coli* before clonal selection and viral sequencing confirmation. Confirmed clones were transfected into HEK 293 cells to produce virions that were then filtered and quantified by p24 ELISA and TZM-bl assay.

### Chemical Inhibitions of Actin Regulators

Actin regulatory proteins were chemically inhibited in migrating primary CD4^+^ T cells to compare their morphologies to HIV-1 infected cells. CDC42 was inhibited by adding 10μM ML 141 (Cayman Chemical, Catolog #18496). Rac proteins (Rac1, Rac1b, Rac2, and Rac3) were inhibited by adding 20μM EHT 1864 (Cayman Chemical Company, Catalog #17258). Pak1/2 were inhibited by adding 50μM IPA-3 (Cayman Chemical Company, Catalog #14759). ARP2/3 was inhibited by adding 200μM CK-666 (Cayman Chemical, Catalog #29038) to the media.

### Transmission Electron Microscopy

Routine transmission electron microscopy (TEM) processing was done as described^65^. In brief, primary human CD4^+^ T cells were negatively selected from freshly isolated PBMC and activated with 25uL anti-CD3 and 25uL anti-CD28 (Ultraleaf purified antibodies) in 10mL RPMI + rhIL-2 (20IU/mL) + IL-15. Cells were kept in a T25 flask and media changes were done every 3-4 days for 22 days. These cells were split into two conditions, Mock infected and HIV-1 infected (ADA-M; TCID50/mL=781,250). Infection was done by spinoculation at 2900 rpm (1965g) for 2 hours prior to plating. 12mL of media supplemented with IL-2 and IL-15 were used to plated CD4^+^ T cells (1*10^6^ cells/mL) onto 100mm diameter cell-culture treated petri dishes that were previously coated with fibronectin. Following 5 days of infection, the cells were gently washed with phosphate-buffered saline and then fixed with 2.5% glutaraldehyde in 0.1 M sodium cacodylate buffer (pH 7.4) at room temperature for 1 hour. The cells were gently scraped off the 100mm tissue culture treated petri dish and pelleted by low-speed centrifugation (100g for 5 minutes). The pellet was fixed for 30 minutes with the same fixative before secondary fixation with 2% osmium tetraoxide on ice for 1 hour. The cells were then stained with 2% uranyl aqueous solution *en bloc* for 1 hour at room temperature, dehydrated with a series of increasing ethanol gradients followed by propylene oxide treatment, and embedded in Embed 812 Resin mixture (Electron Microscopy Sciences). Blocks were cured for 48 h at 65°C and then trimmed into 70 nm ultrathin sections using a diamond knife on a Leica Ultracut 6 and transferred onto 200 mesh copper grids. Sections were counterstained with 1% uranyl acetate in 50% ethanol for 3 minutes at room temperature and in lead citrate for 3 minutes at room temperature, and then examined with a JEOL JSM 1400 transmission electron microscope equipped with two CCD camera for digital image acquisition: Veleta 2K x 2K and Quemesa 11 megapixel (EMSIS, Germany) operated at 100 kV.

### Scanning Electron Microscopy

Scanning electron microscopy (SEM) was done on negatively selected primary human CD4^+^ T cells or cocultures with human myeloid cells grown in RPMI growth media and supplemented with 10% FBS, 20IU/mL IL-2, and antibiotics. Cells were plated on fibronectin-coated coverslips for 5 days. Adherent cells were fixed with 2.5% glutaraldehyde and 1% paraformaldehyde in a 0.12 M sodium cacodylate buffer for 20 minutes at room temperature followed by a fixation for one hour in 1% osmium tetroxide (Electron Microscopy Sciences). The cells were dehydrated through a series of ethyl alcohol/deionized water solutions followed by critical point drying and sputter coating with iridium. Imaging was done using a FEI Teneo LV SEM instrument (Thermo Fisher Scientific).

### Time Lapse Confocal Microscopy

Time lapse confocal imaging was done in an incubated chamber. Image frames were taken at 1 second intervals with either 25X, 40X, or 63X objectives. Videos were generated using a speed rate of 25 frames per second using Zeiss ZenBlue Software. When tracking organelle location using fluorescent dyes, 1μM MitoTracker™ Deep Red FM (ThermoFisher, Catalog number: M22426) was used to detect mitochondria and 1μM LysoTracker™ Blue DND-22 (ThermoFisher, Catalog number: L7525) was used to detect acidic organelles including lysosomes and nuclei.

### Mass Spectrometry

CD4^+^ T cells from 10 HIV-1 negative donors were isolated, activated, and infected with EGFP-containing reporter viruses. 4-7 days post infection, Mock, HIV-1 WT EGFP, and HIV-1 ΔNef EGFP CD4^+^ T cells were suspended at a concentration of 2*10^7^ cells/mL in complete media with IL-2 and IL-15 prior to cell sorting on a BD FACSAria™ II instrument. DAPI staining and mock infected control cells were used to set gates for live and single cells. Cells positive for EGFP in HIV-1 WT EGFP and HIV-1 ΔNef EGFP cultures along with bystander CD4^+^ T cells negative for EGFP were all individually collected. Sorted populations were then washed 3 times in PBS, and cell pellets were stored in -80°C until mass spectrometry was initiated. Cell pellets were resuspended in 9M urea, 50mM Tris, pH8.0. Proteomics analysis of HIV-1 infected and control T cells was done using label-free quantitation at the Weill Cornell Proteomics and Metabolomics Core. Proteins were precipitated using acetone before in-solution trypsin digestion was performed, followed by stage-tip desalting and LC-MS/MS. Each sample was analyzed using a data independent acquisition (DIA) method. The data were searched against customized database containing Uniprot human protein sequences and nine target sequences including HIV-1 proteins and EGFP.

## Acknowledgements

The following reagents were obtained through the NIH HIV Reagent Program, Division of AIDS, NIAID, NIH: HIV-1 ADA (ARP-416), contributed by Dr. Howard Gendelman; HIV-1 92/BR/014 (ARP-1753) and HIV-1 92/BR/004 (ARP-1752), contributed by UNAIDS Network for HIV Isolation and Characterization; and Human rhIL-2 from Dr. Maurice Gately, Hoffmann - La Roche Inc. (ARP-136). Human IL-15 was obtained through the Frederick National Laboratory for Cancer Research Biological Resources Branch pre-clinical repository. The following plasmids were generously provided by Dr. Thorsten Mempel and Dr. Shariq Usmani: HIV-1 (WT) IRES-EGFP (pBR43IeG nef+ R-BaL env), HIV-1 (ΔNef) IRES-EGFP (pBR43IeG nef+ R-BaL env, Nef deleted)^7^. The staff of the Weill Cornell Proteomics and Metabolomics Core carried out the label-free proteome quantitation and the staff of the Weill Cornell Flow Cytometry Core assisted with the cell sorting. This publication resulted in part from research supported by National Institute of Allergy and Infectious Diseases (NIAID) awards UM1 AI126617, co-funded by National Institute on Drug Abuse (NIDA), National Institute of Mental Health (NIMH), and National Institute of Neurological Disorders and Stroke (NINDS).

## Author contributions

RLFO performed the electron and light microscopy, chemical inhibition studies, and genetic screening. DD, JMC, and RLFO performed the mass spectrometry experiments. The conception of the work, the interpretation of results, and manuscript preparation was done by RLFO. All authors reviewed and edited the final manuscript.

### Competing interests

The authors declare no competing financial interests.

Supplementary Information is available for this paper.

### Extended data

**Extended Data Table 1 – Quantitation of label free proteome analysis of HIV-1 infected and control T cells using mass spectrometry**.

## Supplementary information

Supplementary Video 1: Uninfected human CD4^+^ T cell migration on 2-dimensional fibronectin coated dishes (time-lapse, SEM, & TEM).

Supplementary Video 2: HIV-1 infected cells exhibit rounded blebbing prior to apoptosis (time-lapse & TEM).

Supplementary Video 3: Some R5 and R5/X4 viruses induce syncytia formation *in vitro* (time-lapse & TEM).

Supplementary Video 4: Polarized lamellipodial blebbing (time-lapse & TEM).Supplementary Video 5: Needle-like or Rhino phenotype (time-lapse & TEM).

Supplementary Video 6: Abnormal morphologies are unique to HIV-1 infected cells (HIV-1 WT EGFP^+^), (time-lapse).

Supplementary Video 7: Absence of Nef partially but not completely reduces abnormal morphologies. Although Rhino phenotype is not observed, polarized lamellipodial blebbing still occurs, (time-lapse).

Supplementary Video 8: The effects of small molecule inhibitors of actin regulatory proteins on CD4^+^ T cell migration.

Supplementary Video 9: The effects of CK-666, a small molecule inhibitor of ARP2/3, on CD4^+^ T cell migration.

### Data availability

The data presented in this study are available in the supplementary material.

### Human subject data

Deidentified biospecimens were used in these studies. All samples were obtained under institutional IRB approval with documented informed consent.

### Related manuscripts

“A Subset of HIV-1 Controllers Lack Cortical Actin Disruption Indicative of ARP2/3 Inhibition.”

### Preprint servers

bioRxiv

## References

1 Komano, J., Miyauchi, K., Matsuda, Z. & Yamamoto, N. Inhibiting the Arp2/3 complex limits infection of both intracellular mature vaccinia virus and primate lentiviruses. Molecular biology of the cell 15, 5197–5207 (2004). 10.1091/mbc.04-04-0279

2 Welch, M. D. & Way, M. Arp2/3-mediated actin-based motility: a tail of pathogen abuse. Cell host & microbe 14, 242–255 (2013). 10.1016/j.chom.2013.08.011

3 Ospina Stella, A. & Turville, S. All-Round Manipulation of the Actin Cytoskeleton by HIV. Viruses 10 (2018). 10.3390/v10020063

4 Haller, C. et al. The HIV-1 pathogenicity factor Nef interferes with maturation of stimulatory T-lymphocyte contacts by modulation of N-Wasp activity. The Journal of biological chemistry 281, 19618–19630 (2006). 10.1074/jbc.M513802200

5 Lamas-Murua, M. et al. HIV-1 Nef Disrupts CD4(+) T Lymphocyte Polarity, Extravasation, and Homing to Lymph Nodes via Its Nef-Associated Kinase Complex Interface. J Immunol 201, 2731–2743 (2018). 10.4049/jimmunol.1701420

6 Stolp, B. et al. HIV-1 Nef interferes with T-lymphocyte circulation through confined environments in vivo. Proceedings of the National Academy of Sciences of the United States of America 109, 18541–18546 (2012). 10.1073/pnas.1204322109

7 Usmani, S. M. et al. HIV-1 Balances the Fitness Costs and Benefits of Disrupting the Host Cell Actin Cytoskeleton Early after Mucosal Transmission. Cell host & microbe 25, 73–86 e75 (2019). 10.1016/j.chom.2018.12.008

8 Murooka, T. T. et al. HIV-infected T cells are migratory vehicles for viral dissemination. Nature 490, 283–287 (2012). 10.1038/nature11398

9 Stolp, B., Abraham, L., Rudolph, J. M. & Fackler, O. T. Lentiviral Nef proteins utilize PAK2-mediated deregulation of cofilin as a general strategy to interfere with actin remodeling. Journal of virology 84, 3935–3948 (2010). 10.1128/JVI.02467-09

10 Arora, V. K. et al. Lentivirus Nef specifically activates Pak2. Journal of virology 74, 11081–11087 (2000). 10.1128/jvi.74.23.11081-11087.2000

11 Gallego, M. D. et al. WIP and WASP play complementary roles in T cell homing and chemotaxis to SDF-1alpha. International immunology 18, 221–232 (2006). 10.1093/intimm/dxh310

12 Blundell, M. P. et al. Improvement of migratory defects in a murine model of Wiskott-Aldrich syndrome gene therapy. Mol Ther 16, 836–844 (2008). 10.1038/mt.2008.43

13 Lafouresse, F. et al. Wiskott-Aldrich syndrome protein controls antigen-presenting cell-driven CD4+ T-cell motility by regulating adhesion to intercellular adhesion molecule-1. Immunology 137, 183–196 (2012). 10.1111/j.1365-2567.2012.03620.x

14 Krause, M. et al. Fyn-binding protein (Fyb)/SLP-76-associated protein (SLAP), Ena/vasodilator-stimulated phosphoprotein (VASP) proteins and the Arp2/3 complex link T cell receptor (TCR) signaling to the actin cytoskeleton. The Journal of cell biology 149, 181–194 (2000). 10.1083/jcb.149.1.181

15 Sasahara, Y. et al. Mechanism of recruitment of WASP to the immunological synapse and of its activation following TCR ligation. Molecular cell 10, 1269–1281 (2002). 10.1016/s1097-2765(02)00728-1

16 Badour, K., Zhang, J. & Siminovitch, K. A. The Wiskott-Aldrich syndrome protein: forging the link between actin and cell activation. Immunological reviews 192, 98–112 (2003).

17 Badour, K. et al. The Wiskott-Aldrich syndrome protein acts downstream of CD2 and the CD2AP and PSTPIP1 adaptors to promote formation of the immunological synapse. Immunity 18, 141–154 (2003). 10.1016/s1074-7613(02)00516-2

18 Badour, K. et al. Fyn and PTP-PEST-mediated regulation of Wiskott-Aldrich syndrome protein (WASp) tyrosine phosphorylation is required for coupling T cell antigen receptor engagement to WASp effector function and T cell activation. The Journal of experimental medicine 199, 99–112 (2004). 10.1084/jem.20030976

19 Kumari, S. et al. Actin foci facilitate activation of the phospholipase C-gamma in primary T lymphocytes via the WASP pathway. eLife 4 (2015). 10.7554/eLife.04953

20 Janssen, E. et al. A DOCK8-WIP-WASp complex links T cell receptors to the actin cytoskeleton. The Journal of clinical investigation 126, 3837–3851 (2016). 10.1172/JCI85774

21 Krzewski, K., Chen, X., Orange, J. S. & Strominger, J. L. Formation of a WIP-, WASp-, actin-, and myosin IIA-containing multiprotein complex in activated NK cells and its alteration by KIR inhibitory signaling. The Journal of cell biology 173, 121–132 (2006). 10.1083/jcb.200509076

22 Pauker, M. H. et al. WASp family verprolin-homologous protein-2 (WAVE2) and Wiskott-Aldrich syndrome protein (WASp) engage in distinct downstream signaling interactions at the T cell antigen receptor site. The Journal of biological chemistry 289, 34503–34519 (2014). 10.1074/jbc.M114.591685

23 Choe, E. Y., Schoenberger, E. S., Groopman, J. E. & Park, I. W. HIV Nef inhibits T cell migration. The Journal of biological chemistry 277, 46079–46084 (2002). 10.1074/jbc.M204698200

24 Park, I. W. & He, J. J. HIV-1 Nef-mediated inhibition of T cell migration and its molecular determinants. Journal of leukocyte biology 86, 1171–1178 (2009). 10.1189/jlb.0409261

25 Janardhan, A., Swigut, T., Hill, B., Myers, M. P. & Skowronski, J. HIV-1 Nef binds the DOCK2-ELMO1 complex to activate rac and inhibit lymphocyte chemotaxis. PLoS biology 2, E6 (2004). 10.1371/journal.pbio.0020006

26 Dyer, W. B. et al. Lymphoproliferative immune function in the Sydney Blood Bank Cohort, infected with natural nef/long terminal repeat mutants, and in other long-term survivors of transfusion-acquired HIV-1 infection. Aids 11, 1565–1574 (1997).

27 Kestler, H. W., 3rd et al. Importance of the nef gene for maintenance of high virus loads and for development of AIDS. Cell 65, 651–662 (1991).

28 Lindemann, D. et al. Severe immunodeficiency associated with a human immunodeficiency virus 1 NEF/3’-long terminal repeat transgene. The Journal of experimental medicine 179, 797–807 (1994). 10.1084/jem.179.3.797

29 Hanna, Z. et al. Nef harbors a major determinant of pathogenicity for an AIDS-like disease induced by HIV-1 in transgenic mice. Cell 95, 163–175 (1998). 10.1016/s0092-8674(00)81748-1

30 Thomas, A. et al. Involvement of the Rac1-IRSp53-Wave2-Arp2/3 Signaling Pathway in HIV-1 Gag Particle Release in CD4 T Cells. Journal of virology 89, 8162–8181 (2015). 10.1128/JVI.00469-15

31 Aggarwal, A., Stella, A. O., Henry, C. C., Narayan, K. & Turville, S. G. Embedding of HIV Egress within Cortical F-Actin. Pathogens 11 (2022). 10.3390/pathogens11010056

32 Deacon, N. J. et al. Genomic structure of an attenuated quasi species of HIV-1 from a blood transfusion donor and recipients. Science 270, 988–991 (1995).

33 Learmont, J. et al. Long-term symptomless HIV-1 infection in recipients of blood products from a single donor. Lancet 340, 863–867 (1992).

34 Zhang, Q. et al. DOCK8 regulates lymphocyte shape integrity for skin antiviral immunity. The Journal of experimental medicine 211, 2549–2566 (2014). 10.1084/jem.20141307

35 Bergert, M., Chandradoss, S. D., Desai, R. A. & Paluch, E. Cell mechanics control rapid transitions between blebs and lamellipodia during migration. Proceedings of the National Academy of Sciences of the United States of America 109, 14434–14439 (2012). 10.1073/pnas.1207968109

36 Derivery, E. et al. Free Brick1 is a trimeric precursor in the assembly of a functional wave complex. PloS one 3, e2462 (2008). 10.1371/journal.pone.0002462

37 Davidson, A. J., Amato, C., Thomason, P. A. & Insall, R. H. WASP family proteins and formins compete in pseudopod-and bleb-based migration. The Journal of cell biology 217, 701–714 (2018). 10.1083/jcb.201705160

38 Steffen, A. et al. Sra-1 and Nap1 link Rac to actin assembly driving lamellipodia formation. The EMBO journal 23, 749–759 (2004). 10.1038/sj.emboj.7600084

39 Obeidy, P. et al. Partial loss of actin nucleator actin-related protein 2/3 activity triggers blebbing in primary T lymphocytes. Immunol Cell Biol 98, 93–113 (2020). 10.1111/imcb.12304

40 Cook, S. A. et al. HEM1 deficiency disrupts mTORC2 and F-actin control in inherited immunodysregulatory disease. Science 369, 202–207 (2020). 10.1126/science.aay5663

41 Han, Y. et al. Formin-like 1 (FMNL1) is regulated by N-terminal myristoylation and induces polarized membrane blebbing. The Journal of biological chemistry 284, 33409–33417 (2009). 10.1074/jbc.M109.060699

42 Rotty, J. D. et al. Arp2/3 Complex Is Required for Macrophage Integrin Functions but Is Dispensable for FcR Phagocytosis and In Vivo Motility. Developmental cell 42, 498–513 e496 (2017). 10.1016/j.devcel.2017.08.003

43 Kumar, S. et al. Cdc42 regulates neutrophil migration via crosstalk between WASp, CD11b, and microtubules. Blood 120, 3563–3574 (2012). 10.1182/blood-2012-04-426981

44 Fritz-Laylin, L. K., Lord, S. J. & Mullins, R. D. WASP and SCAR are evolutionarily conserved in actin-filled pseudopod-based motility. J Cell Biol 216, 1673–1688 (2017). 10.1083/jcb.201701074

45 Leithner, A. et al. Diversified actin protrusions promote environmental exploration but are dispensable for locomotion of leukocytes. Nature cell biology 18, 1253–1259 (2016). 10.1038/ncb3426

46 Chikina, A. S., Svitkina, T. M. & Alexandrova, A. Y. Time-resolved ultrastructure of the cortical actin cytoskeleton in dynamic membrane blebs. The Journal of cell biology 218, 445–454 (2019). 10.1083/jcb.201806075

47 Henson, J. H. et al. Arp2/3 complex inhibition radically alters lamellipodial actin architecture, suspended cell shape, and the cell spreading process. Molecular biology of the cell 26, 887–900 (2015). 10.1091/mbc.E14-07-1244

48 Chanez-Paredes, S., Montoya-Garcia, A. & Schnoor, M. Cellular and pathophysiological consequences of Arp2/3 complex inhibition: role of inhibitory proteins and pharmacological compounds. Cell Mol Life Sci 76, 3349–3361 (2019). 10.1007/s00018-019-03128-y

49 Sokolova, O. S. et al. Structural Basis of Arp2/3 Complex Inhibition by GMF, Coronin, and Arpin. Journal of molecular biology 429, 237–248 (2017). 10.1016/j.jmb.2016.11.030

50 Weaver, A. M., Young, M. E., Lee, W.-L., & Cooper, J. A. Integration of signals to the Arp2/3 complex. Current opinion in cell biology 15, 23–30 (2003). 10.1016/s0955-0674(02)00015-7

51 Spear, M. et al. HIV-1 triggers WAVE2 phosphorylation in primary CD4 T cells and macrophages, mediating Arp2/3-dependent nuclear migration. The Journal of biological chemistry 289, 6949–6959 (2014). 10.1074/jbc.M113.492132

52 Usmani, S. M. & Mempel, T. R. Cancer cells relax and resist cytotoxic attack. Immunity 54, 853–855 (2021). 10.1016/j.immuni.2021.04.017

53 Tello-Lafoz, M. et al. Cytotoxic lymphocytes target characteristic biophysical vulnerabilities in cancer. Immunity 54, 1037–1054 e1037 (2021). 10.1016/j.immuni.2021.02.020

54 Wiskott, A. Familiärer, angeborener Morbus werlhoffi? Monatsschr. Kinderh. 68 (1937).

55 Aldrich, R. A., Steinberg, A. G. & Campbell, D. C. Pedigree demonstrating a sex-linked recessive condition characterized by draining ears, eczematoid dermatitis and bloody diarrhea. Pediatrics 13, 133–139 (1954).

56 Cooper, M. D., Chae, H. P., Lowman, J. T., Krivit, W. & Good, R. A. Wiskott-Aldrich syndrome. An immunologic deficiency disease involving the afferent limb of immunity. Am J Med 44, 499–513 (1968). 10.1016/0002-9343(68)90051-x

57 Somech, R. et al. Disruption of Thrombocyte and T Lymphocyte Development by a Mutation in ARPC1B. J Immunol 199, 4036–4045 (2017). 10.4049/jimmunol.1700460

58 Brigida, I. et al. T-cell defects in patients with ARPC1B germline mutations account for combined immunodeficiency. Blood 132, 2362–2374 (2018). 10.1182/blood-2018-07-863431

59 Sindram, E. et al. ARPC5 deficiency leads to severe early-onset systemic inflammation and mortality. Disease models & mechanisms 16 (2023). 10.1242/dmm.050145

60 Alkhairy, O. K. et al. RAC2 loss-of-function mutation in 2 siblings with characteristics of common variable immunodeficiency. J Allergy Clin Immunol 135, 1380–1384 e1381-1385 (2015). 10.1016/j.jaci.2014.10.039

61 Tangye, S. G. et al. Human inborn errors of the actin cytoskeleton affecting immunity: way beyond WAS and WIP. Immunol Cell Biol 97, 389–402 (2019). 10.1111/imcb.12243

62 Castro, C. N. et al. NCKAP1L defects lead to a novel syndrome combining immunodeficiency, lymphoproliferation, and hyperinflammation. The Journal of experimental medicine 217 (2020). 10.1084/jem.20192275

63 Cook, S., Lenardo, M. J. & Freeman, A. F. HEM1 Actin Immunodysregulatory Disorder: Genotypes, Phenotypes, and Future Directions. J Clin Immunol (2022). 10.1007/s10875-022-01327-0

64 Liu, M. et al. WAVE2 suppresses mTOR activation to maintain T cell homeostasis and prevent autoimmunity. Science 371 (2021). 10.1126/science.aaz4544

65 Ikegame, S. et al. Metagenomics-enabled reverse-genetics assembly and characterization of myotis bat morbillivirus. Nat Microbiol (2023). 10.1038/s41564-023-01380-4

